# FabElm_BarcodeDb: *matK* barcode database of legumes

**DOI:** 10.1101/241703

**Authors:** Bharat Kumar Mishra, Sakshi Chaudhary, Jeshima Khan Yasin

**Affiliations:** Division of Genomic Resources, ICAR-NBPGR, PUSA campus, New Delhi, India- 110012

**Keywords:** DNA Barcoding, Leguminosae, matK, splicing, phylogeny, diversity, molecular clock

## Abstract

**Background:** DNA barcoding is an imperative implementation of chloroplast *rbcL* and *matK* regions exploited as standard molecular barcodes for species identification. *MatK* is highly conserved in plants and has been used extensively as a phylogenetic marker for classification of plants. In this study *matK* sequences of Leguminosae were retrieved for variant analysis and phylogentics. From online resources, maturase sequences were retrieved; redundant sequences and partials along with poor quality reads were filtered to compile 3639 complete non-redundant *matK* sequences and constructed into a database for ready reference. The database FabElm_BarcodeDb made available at app.bioelm.com was constructed using available sequence resources.

**Results:** The chloroplast genome of plants contains *matK* gene of 1500 bp, positioned between intron of *trnK* associated in-group II intron splicing. Mitochondrial *matR* and genomic *matN* sequences were compared with chloroplast *matK*. These maturase sequences share regions of homology with chloroplast and mitochondrial regions and are expected to be regulated by miRNA in producing splice variants contributing to speciation.

**Conclusion:** Base substitution rates of nuclear maturase were comparable with mitochondrial maturase and are different from *matK* sequences. Hence, few identified species in this investigation were clustered with other tribes when analysed using *matK. MatK* is effective in resolving the species level variations as splicing contributes to speciation; but utilization of *matK* alone as a barcode marker for legumes is dubious, as it could not resolve some species identity.

**Abbreviations:**

InDels: Insertions and deletions
ITS: Internal transcribed spacer
*matK*: Maturase K
*rbcL*: Ribulose-bisphosphate carboxylase gene

## Background

Leguminosae, the third largest family of angiosperms representing 730 genera and beyond 19,400 species of which living ones are 16371 across globe (Mabberley, 1997 and Lewis et al., 2005). Legumes are of wide economic significance as crops of food, fodder, fibre, timber, fuel, oils, chemicals and medicines contributing to food and nutritional security (Ramya et al., 2013; Yasin, 2015; Yasin, 2016). Legumes stretch in habitat from annual herbs to huge trees and are pleasingly connoted all over tropical and temperate regions around the globe (Rundel, 1989). Legumes fix atmospheric nitrogen through root-nodulating symbiotic bacteria bringing in sustainability to encounter their metabolic nitrogen demands. Entire legumes contributes to the terrestrial nitrogen cycle regardless of their nodule forming carbon footprint (Sprent, 1994; Sprent, 2001).

The sequence variations investigation at genus level revealed extensively studied nuclear, chloroplast and mitochondrial specific genes; rbcL, matK and trnK by researchers for plant systematics (Neuhaus and Link, 1987). Significantly, the genes who leads the chloroplast genome are classified in two different classes; one class codes for the components of photosynthetic apparatus while, the other class codes for their gene expression and regulation systems. A protein convoluted in the splicing process is encoded by a gene prominently known as maturase K *(matK)*.

The continuous chloroplast inheritance in tens of thousands of plant is controlled by *matK* gene (Turmel et al., 2006) as a typical bio-molecular marker for phylogenetic studies and applications (CBOL, 2009). The absence of *matK* gene in parasites of Cuscuta (Funk et al., 2007; McNeal et al., 2009) and Orchid (Delannoy et al., 2011) genus results in the loss of *matK* associated group II introns, making *matK* gene indispensable for group II intron splicing. Minute information essential in the direction of probable regulator to specific splicing factor can be studied by expression characterisation of *matK* or/and their intron targets.

*MatR* sequences are highly conserved and well represented across angiosperm lineages (Adams et al., 2002) anticipating an important introns splicing factor in mitochondria. Furthermore, RNA-editing measures to reinstate the highly conserved amino acids across species (Stern et al, 2010), added with *matR* coding a functional mitochondrial protein. Straight evidences are unavailable to illustrate the splicing actions of *matR* being a functional protein. The earlier studies intricate *matR* relationship with many pre-RNAs *in-vivo*, fitting with group II introns (Schmitz et al, 2005 and Grewe et al, 2014), indicating *matR* as a substantial splice factor comparable with *matK* (Sultan at el., 2016).

Phylogenetic reconstruction of Leguminosae is critical to understand the source and divergence of these organically and frugally significant angiosperms. Widespread phylogeny reports of Leguminosae started with plastid gene, *rbcL* (Chase et al., 1993; Doyle, 1987; Doyle, 1997) followed by *matK* (Wojciechowski et al., 2004). Our study is directed towards evaluating the efficiency of *matK* as solitary biomarker and the probable impact of coevolved splicing in comparisons with other maturase genes.

## Construction, content and data analyses

### Taxon sampling

Complete set of available *matK* sequences including 235 legume genera documented by Polhill (1994) were incorporated in this study. Available chloroplast *matK* sequences from 730 genera were included for further analyses, delegating 19,200 species of legumes. Suitable out-group taxa from Quillajaceae, Polygalaceae, Surianaceae, Rosaceae from Fabales and Rosales (Vauquelinia) were selected based on previous molecular phylogenetic reports of eurosids/ fabids (Soltis et al., 2000; Soltis et al., 1995; Kajita et al., 2001; Steele and Vilgalys, 1994; and Persson, 2001).

### DNA sequence data collection

The complete set of reported Leguminosae species list were retrieved from taxonomy database of National Centre for Biotechnology information (NCBI). A total of 10337 nucleotide sequences and popset were retrieved from NCBI. Redundant sequences and partials along with poor quality reads were filtered to compile 3639 complete non-redundant *matK* sequences.

### Multiple sequence analysis *matK*

The *matK* sequences were primarily mounted up into a data file, curated then aligned implementing Multiple Sequence Comparison by Log-Expectation; MUSCLE (Edgar, 2004) with benchmark pair-wise and multiple sequence alignment parameters as default settings.

### *In-silico* prediction of miRNAs targeting *matK* sequences

PsRNAtarget (Dai and Zhao, 2011) was utilized to predict the miR target regions of maturase. The reported miRBase viridiplantae miRNAs were used to findout the miRNAs targeting maturase gene sequences.

### Domain and splice junction prediction

A set of 13 *Medicago spp*. nuclear maturase sequences were subjected to splice junction predictions. Three sequences were filtered out as the reported sequence length was insufficient for analyses (<200 b). Acceptor and donor sites were plotted and probable junctions from both direct and complementary strands were marked.

### Co-expression network analysis

Maturase genes and their putative co-expression members were identified in gene expression and splice variant regulation through unweighted network analyses. Outlier members were eliminated to redraw the network with linked genes.

### Phylogenetic analyses

The sequences were analysed by Minimum Evolution Tree method through MEGA7 (Kumar et al., 2016). Statistical parameters were set to default for universal phylogenetic reconstruction. Additionally, the evolutionary clock time for *matK* sequences of different legumes was estimated by MEGA7.

### Results and Discussion

With an objective to correlate speciation to phylogenetic relationship among Leguminosae using chloroplast *matK* gene sequences accessible from free resources and knowledge base the present investigation was carried out and constructed into a database. Of the total expected >19,000 *matK* sequences for known species of Fabaceae, only 3639 complete nonredundant *matK* sequences were available from the database indicating the larger gap in available sequence resources.

### Characteristics of the *matK* sequences

*MatK* gene in legumes length stretches from 1476 nucleotide bases corresponds to 492 numbers of amino acids in *Erythrostemo gilliesii* to 1545 nucleotide bases corresponds to 515 numbers of amino acids in more than a few dalbergioid taxa species; Adesmia and Pictetia. Our absolute *matK* data-set comprises 1674 aligned positions corroborating 1042 putative parsimony-dependent characters representing 73% of total nucleotide bases excluding 244 InDels and absent characters amid analysed 330 taxa. Among 552420 characters representing the complete data-set, 1.3% are missing while 14.6% corresponding 80500 bases are InDels and other left out positions. Comparatively, the previous comprehensive study of *rbcL* in legume (Kajita et al., 2001) comprising sequences of 1404 nucleotide bases in length alignments without/few InDels having 530 informative characters among the 319 sequences which are significantly less. Maturase sequence of chloroplast is less informative when compared to *matR* of mitrochondria.

*MatK* sequences are enriched with *matK* and type II intron maturase domains (fig. 1) whereas, the *matR* comprises RT like super family and RT_G2 intron domains. Nuclear maturase are with matK-N super family domains as well as intron maturase regions. Additional conserved domains are predicted and depicted in (Fig. 2). The position and length of domains varies among the sequences as well as between chloroplast, mitochondria and nuclear maturase genes (Fig. 1–3). There are sequences of same species with significant domain variations indicating chances of getting it identified by mistake or there could be subspecies level variations handled with insufficient markers or names (Fig. 3).

**Figure 1.**
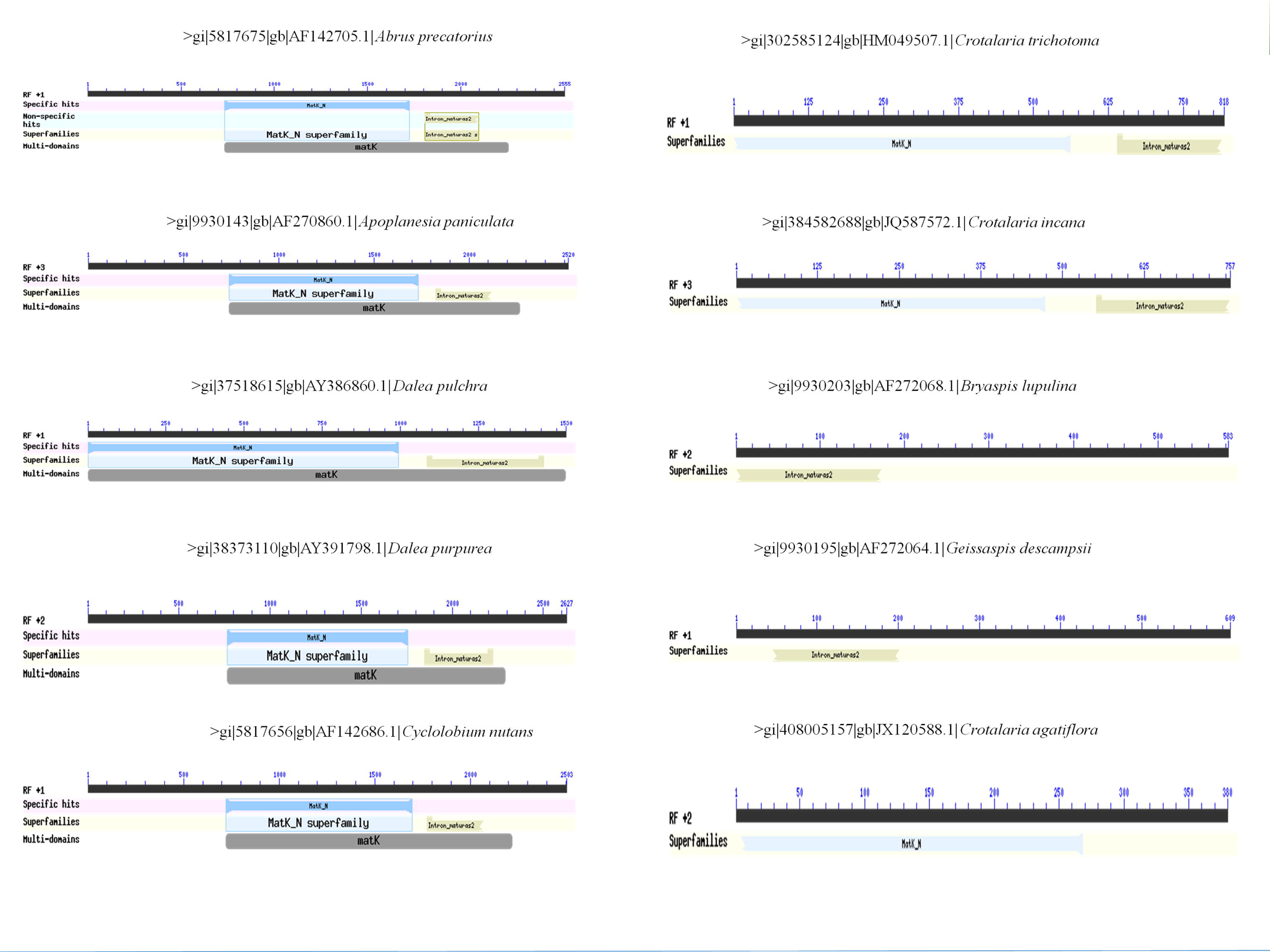
MatK intron domains.

**Figure 2.**
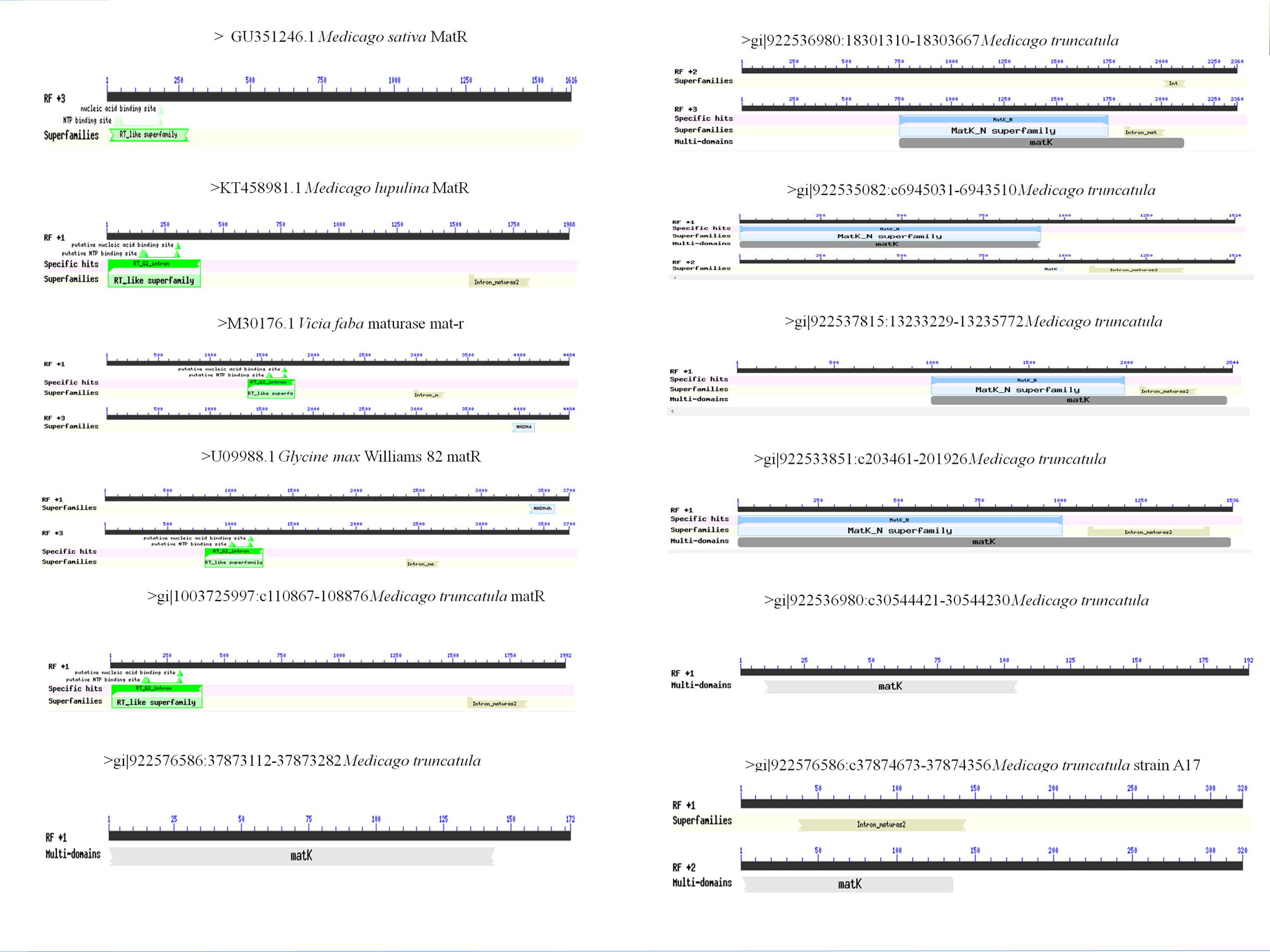
Conserved domains.

**Figure 3.**
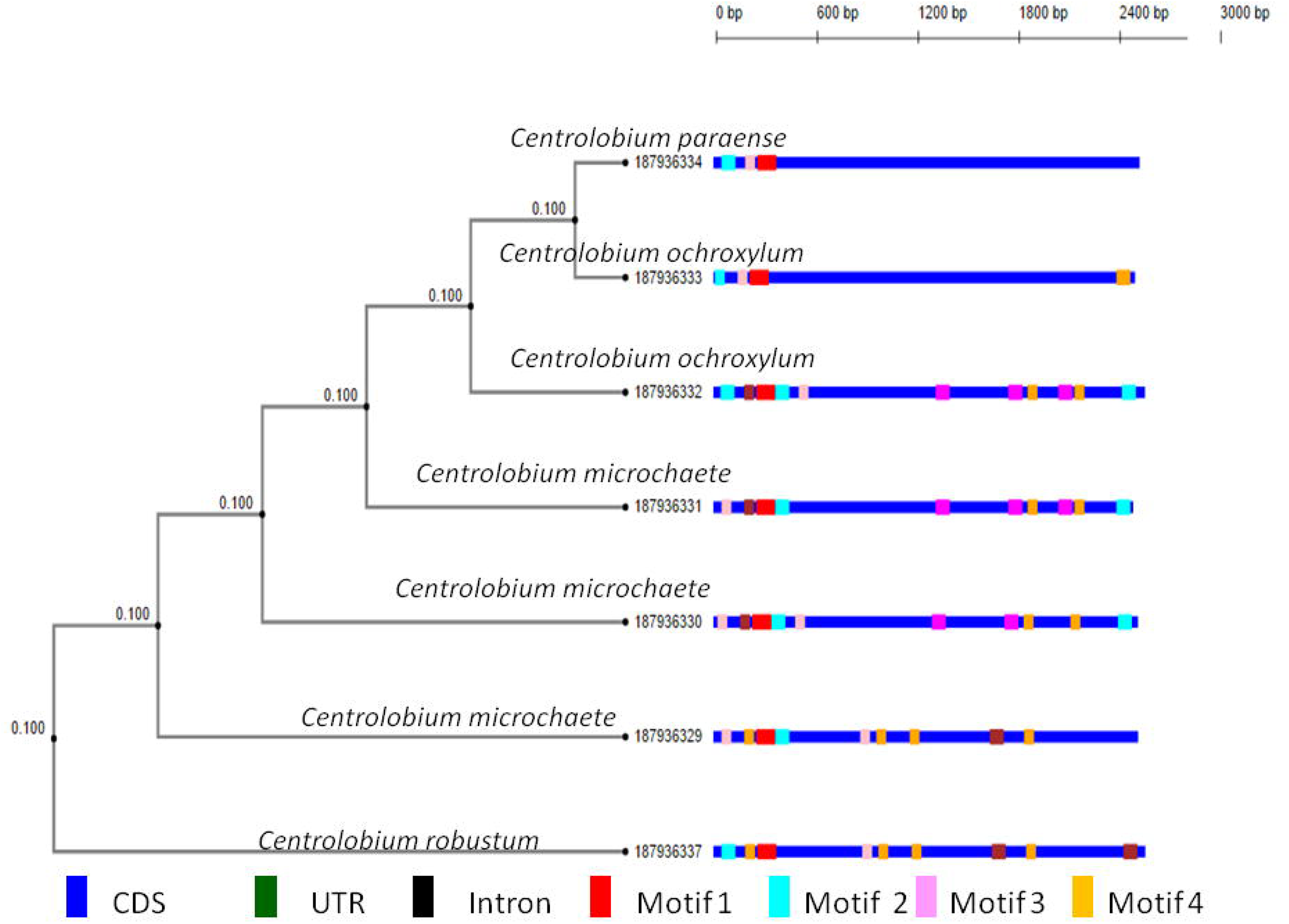
Motif diversity.

### Phylogenetic reconstruction

The maximum parsimony (MP) tree with strict consensus is extremely resolved and illustrated in Figs. 4–5. Semi-strict consensus MP trees determine three nodes. Interestingly, 50% majority-rule tree resulted in seven fully unresolved nodes. Deviations in grouping occurred among major genus where are clustered with diverse genus (Fig.6–24).

**Figure 4.**
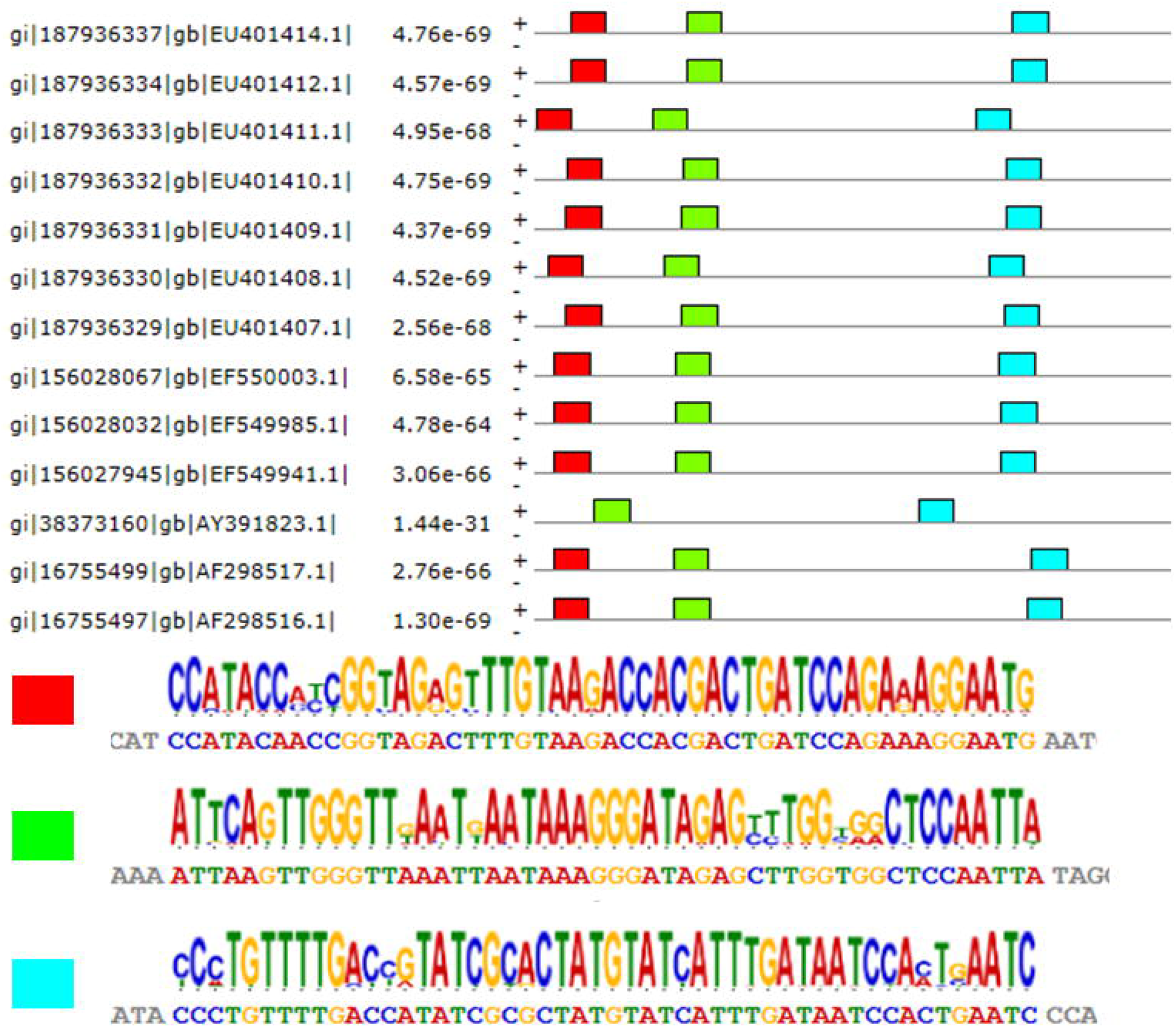
Novel motif.

**Figure 5.**
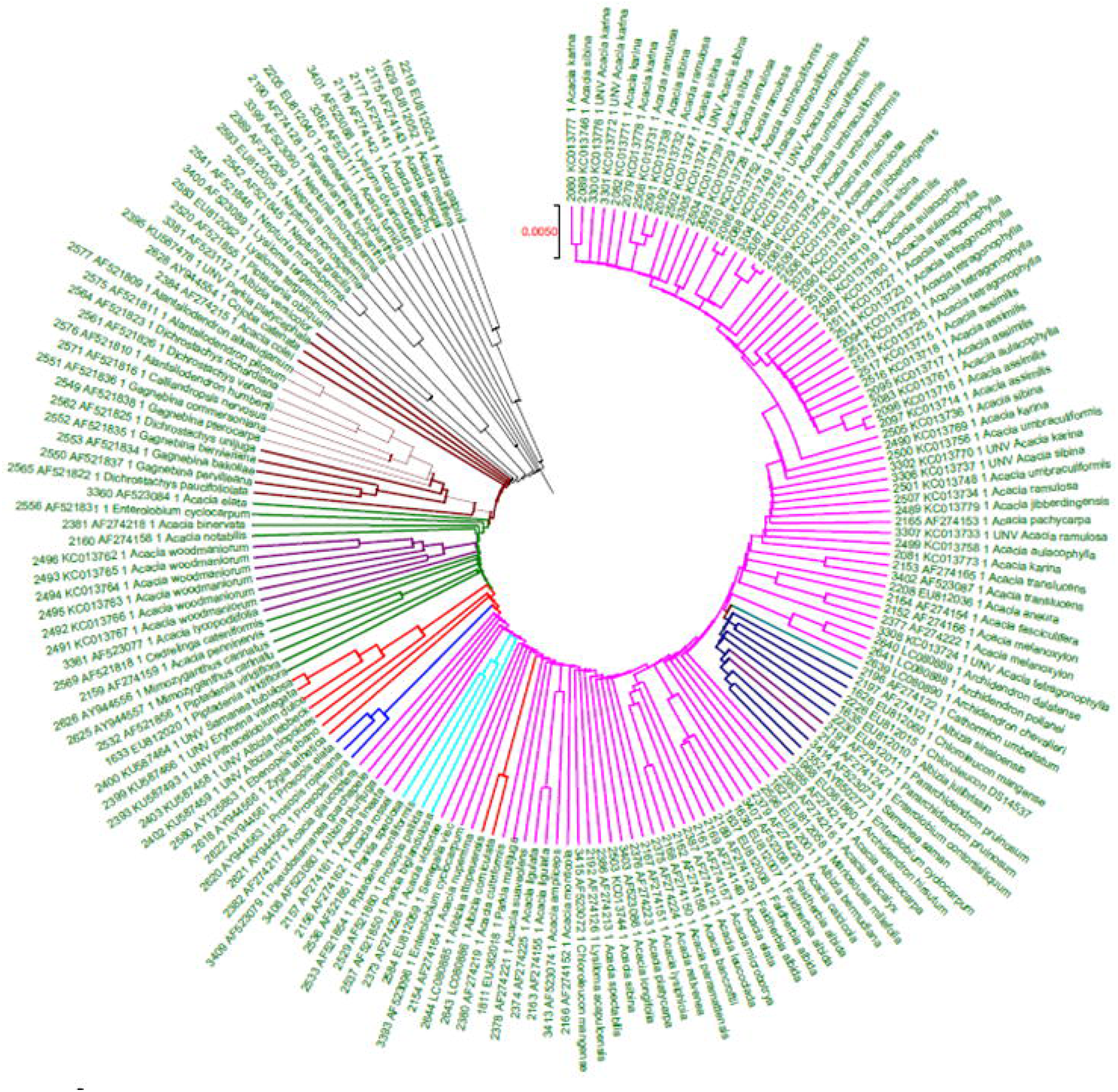
Acacia group.

**Figure 6.**
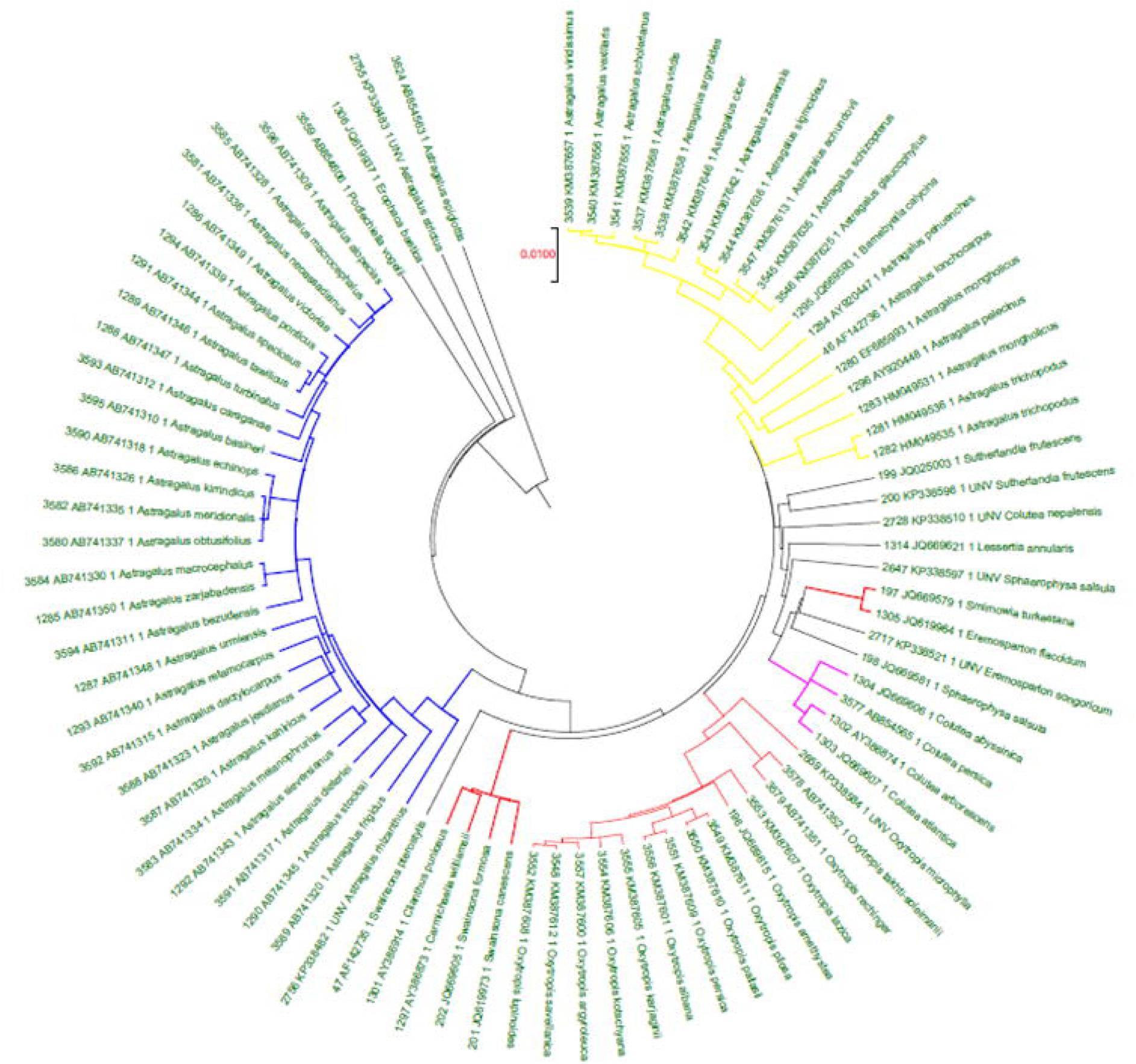
Astragalus group.

**Figure 7.**
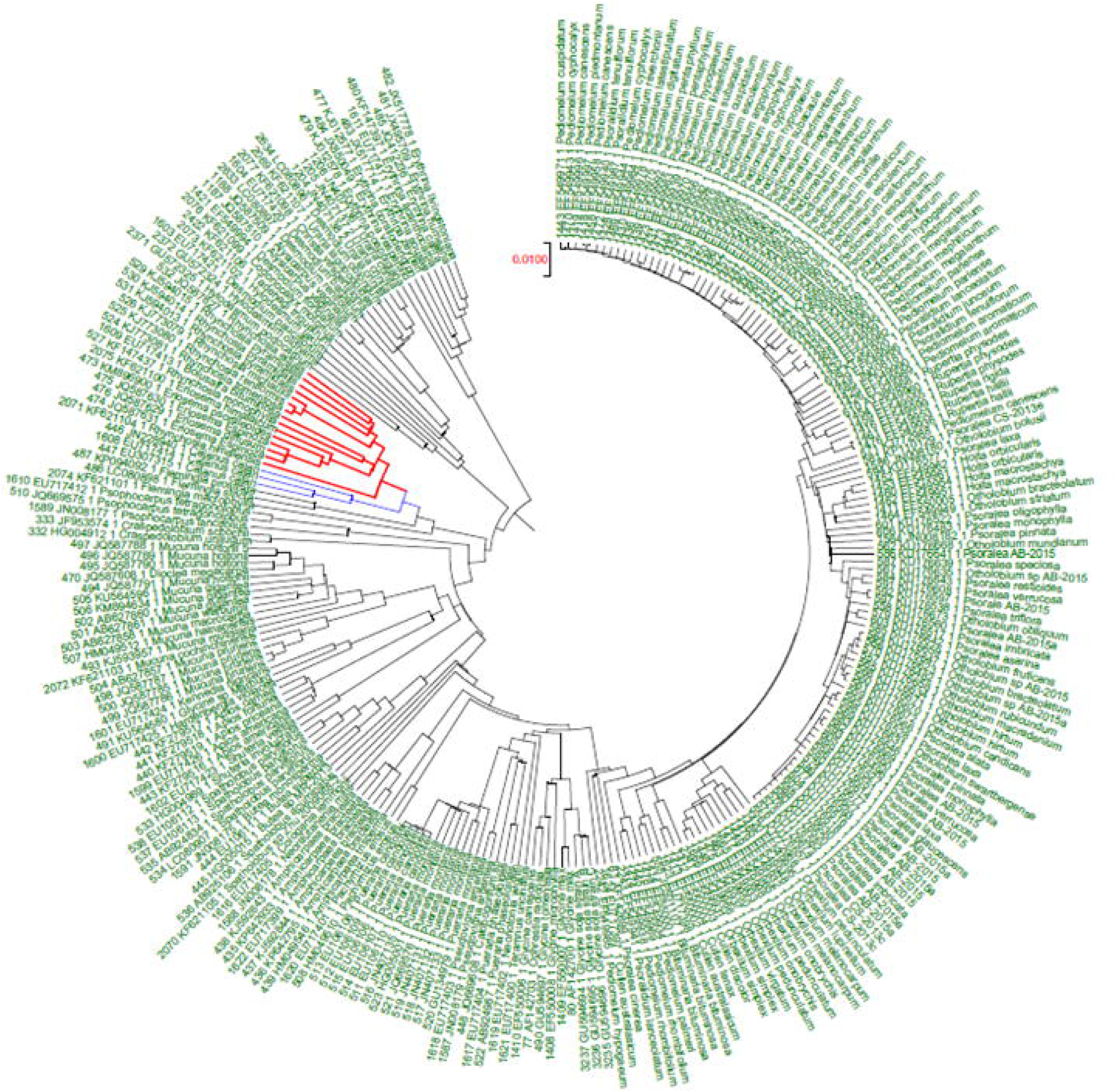
Cajanus group.

**Figure 8.**
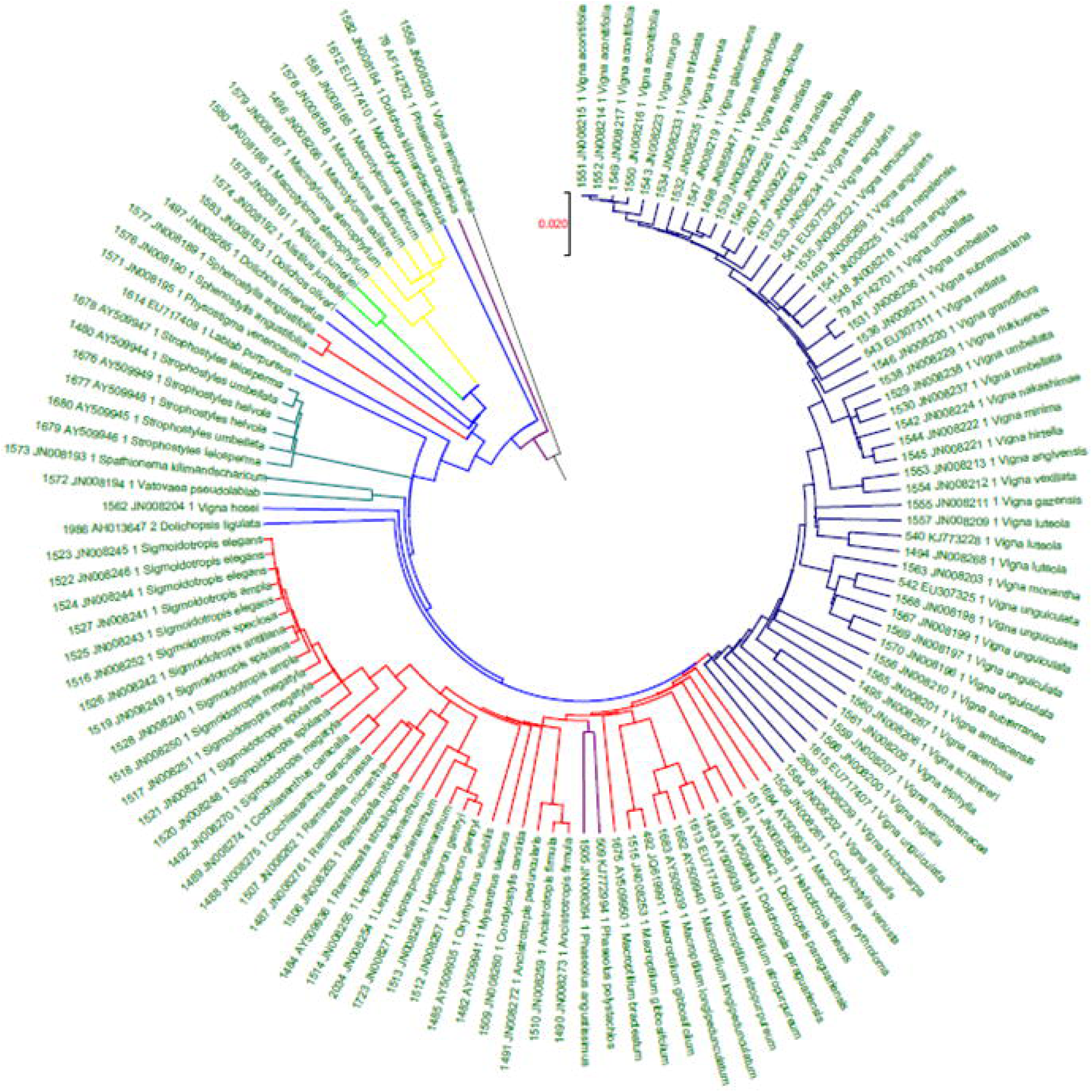
Dolichos group.

**Figure 9.**
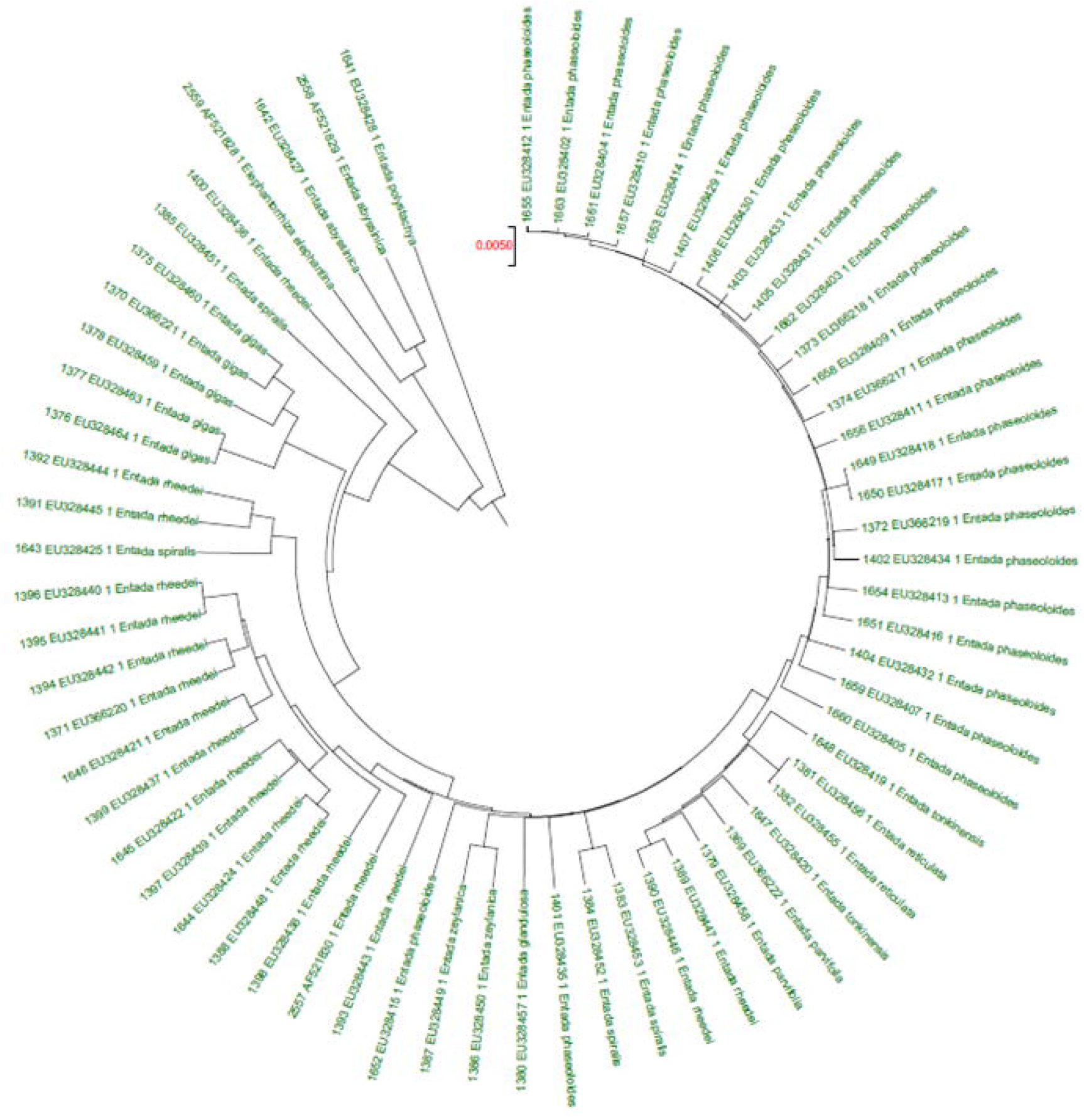
Entada group.

**Figure 10.**
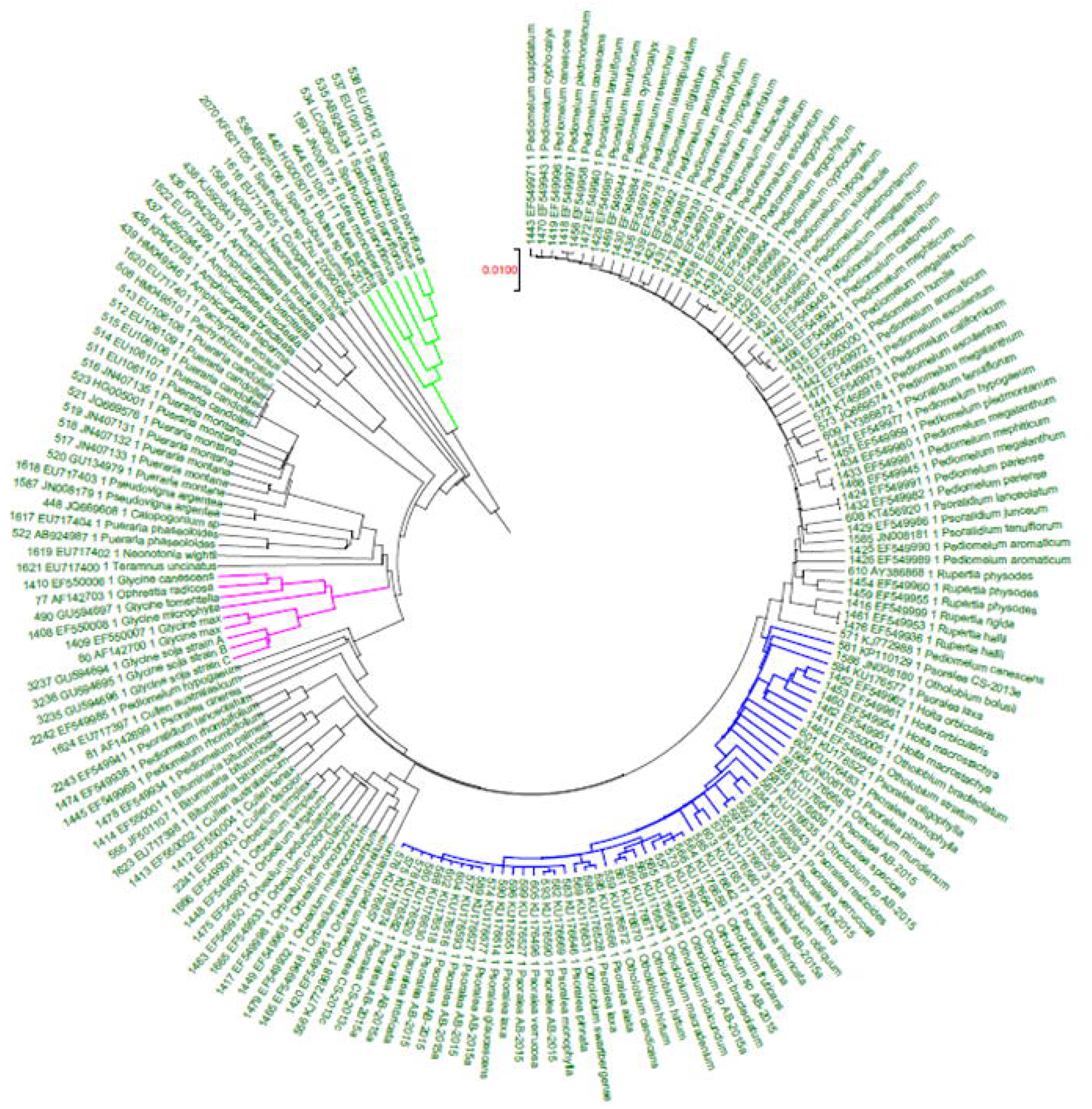
Glycine group.

**Figure 11.**
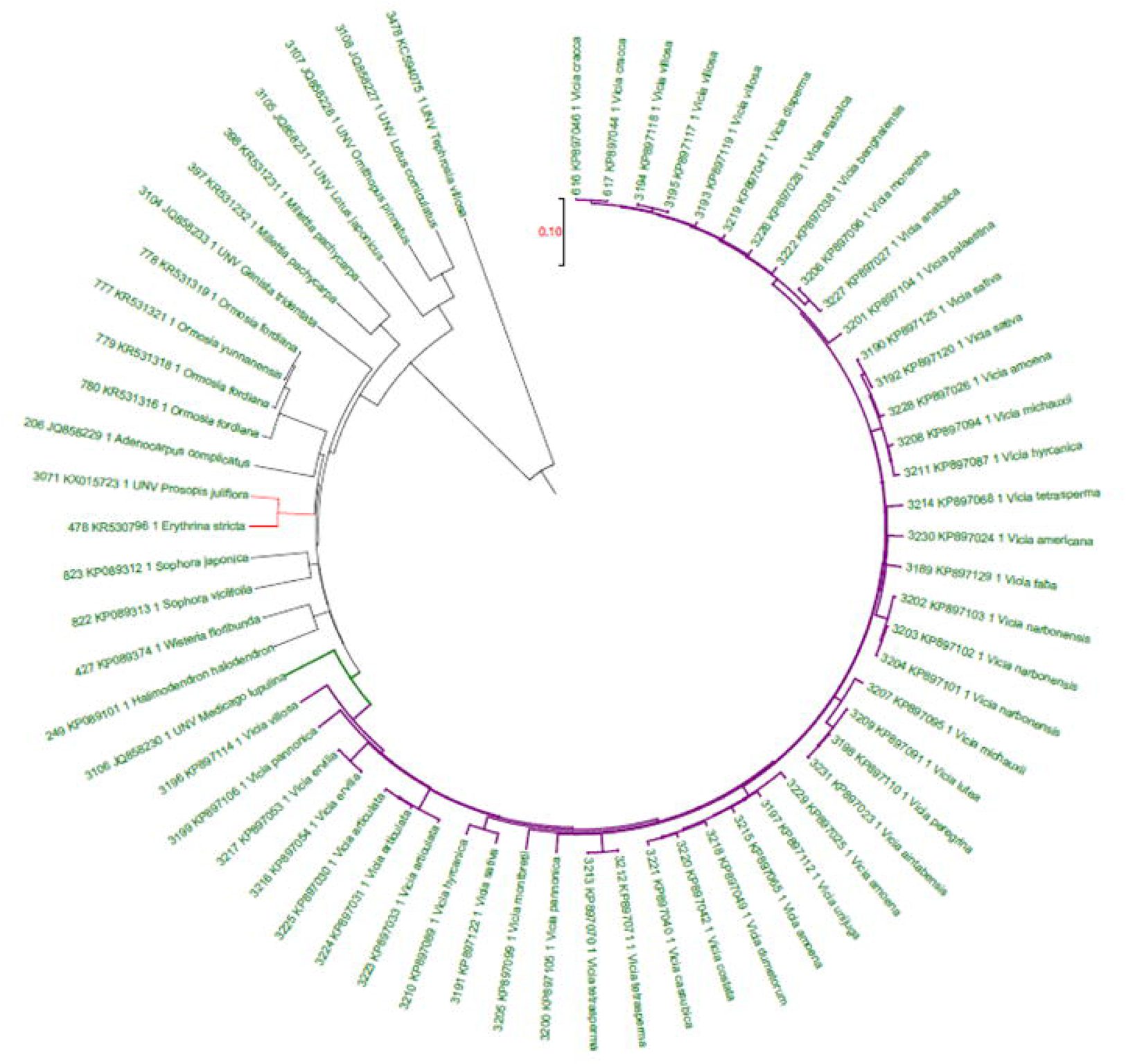
Medicago and vicia group.

**Figure 12.**
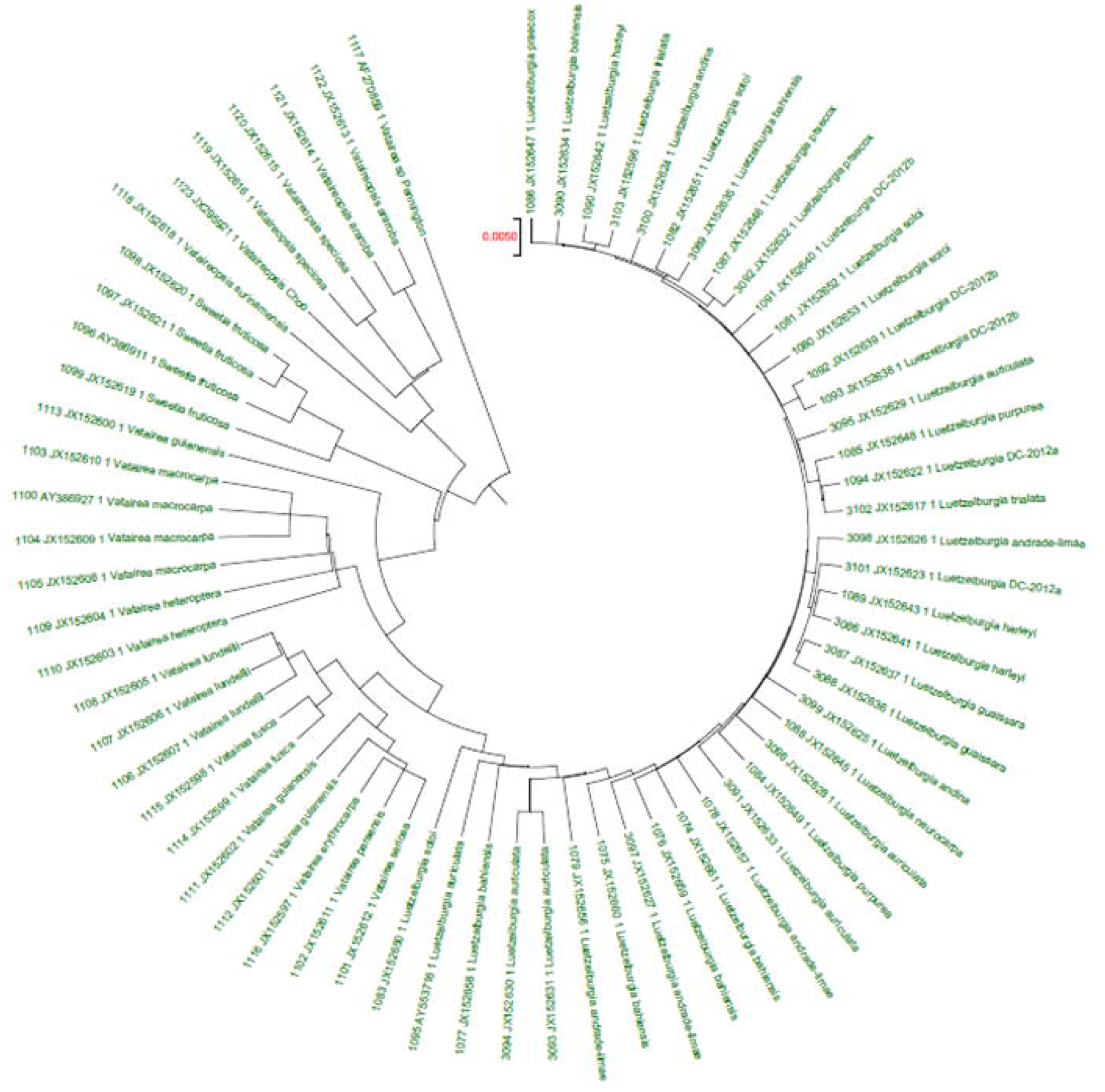
Sweetia group.

**Figure 13.**
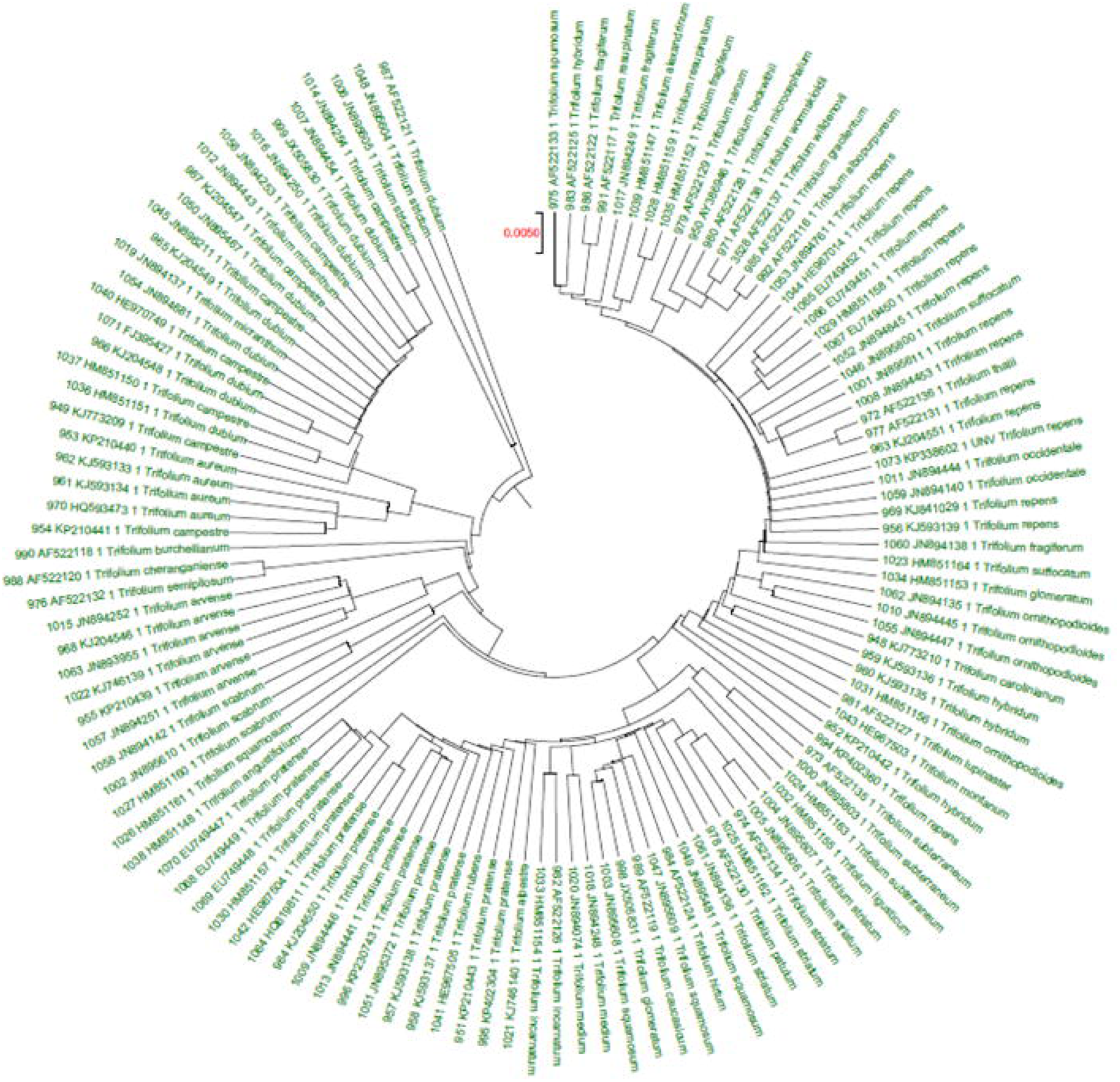
Trifolium group.

**Figure 14.**
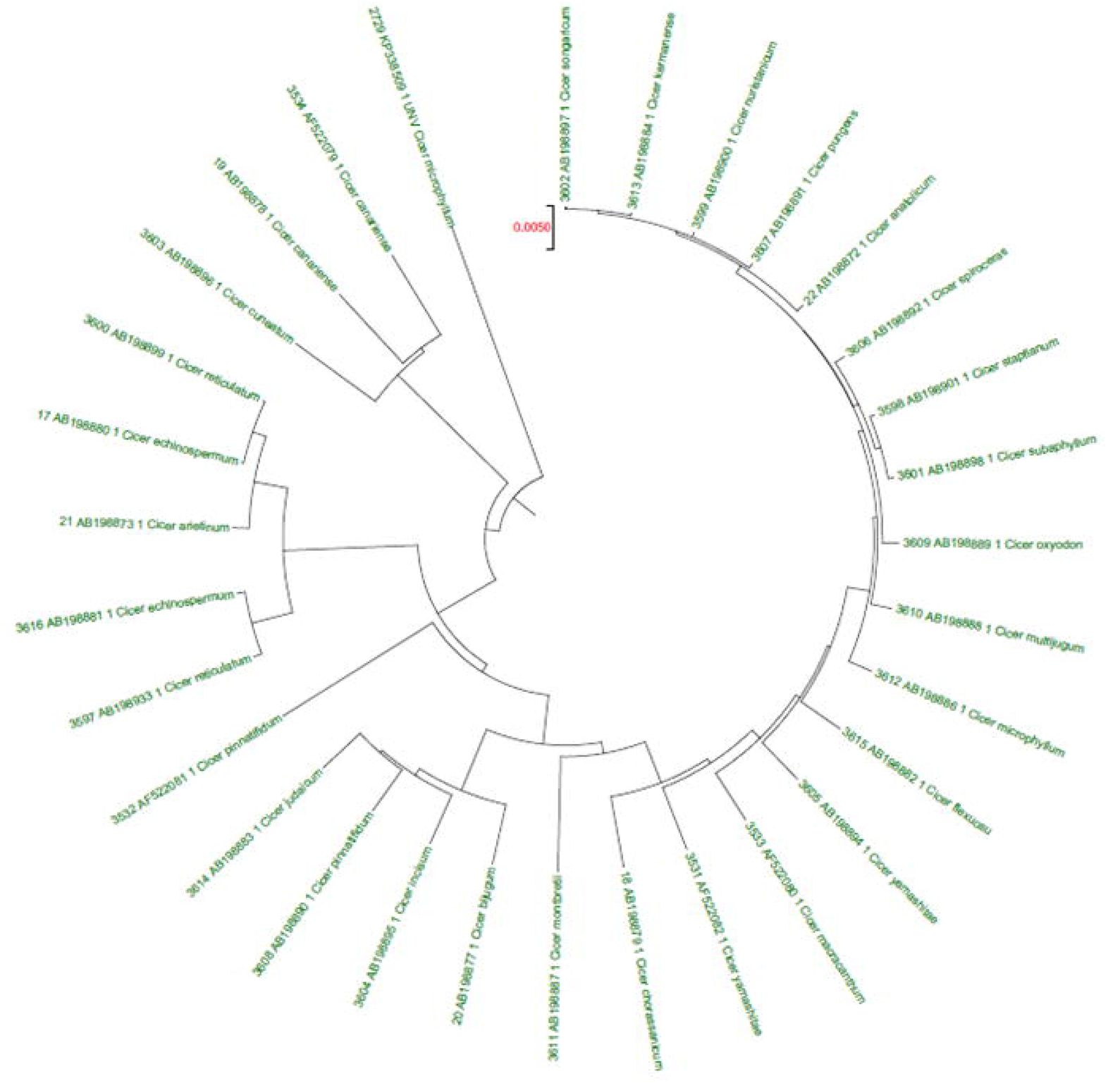
Medicago with trifolium.

**Figure 15.**
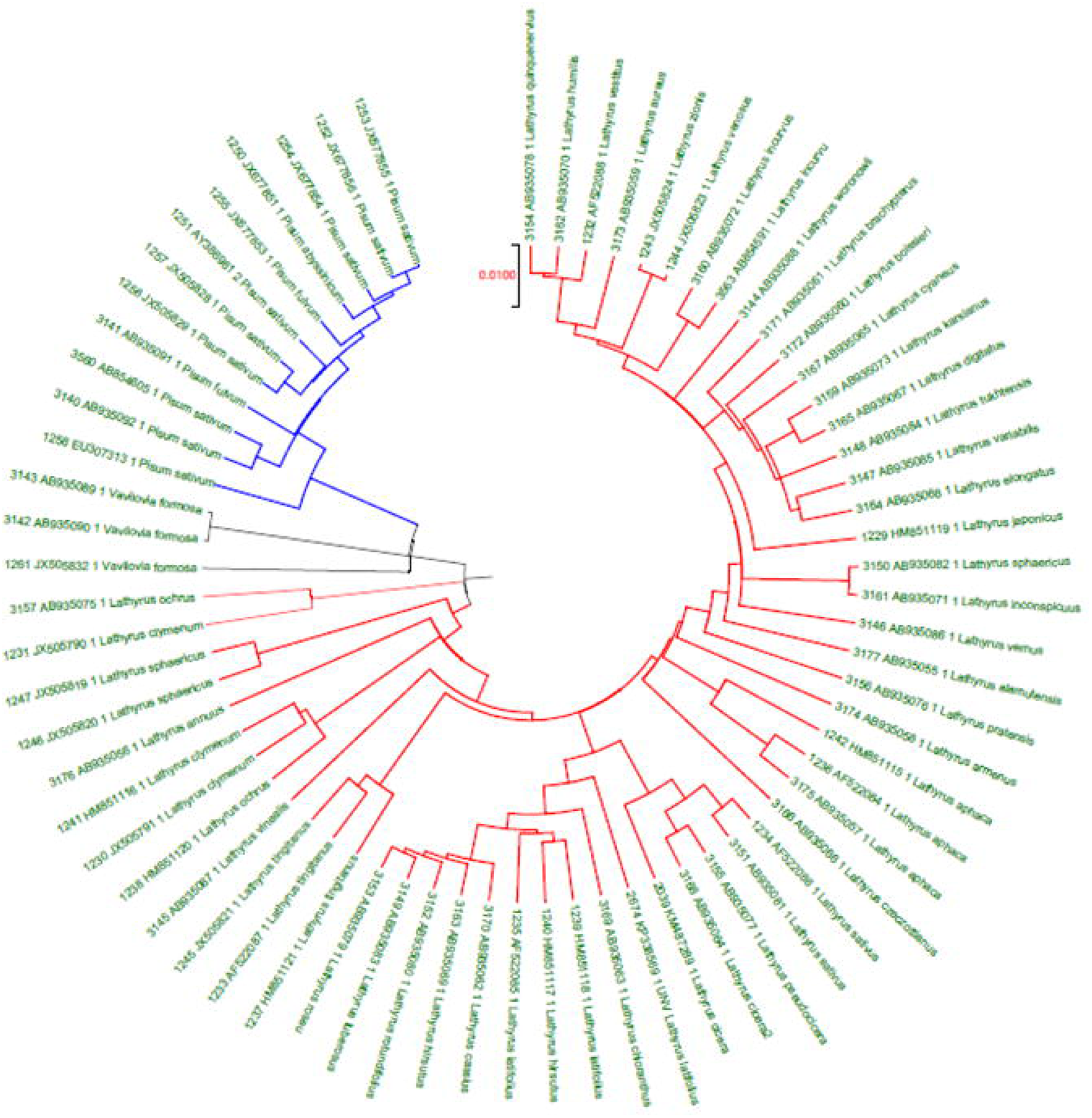
Lathyrus group.

**Figure 16.**
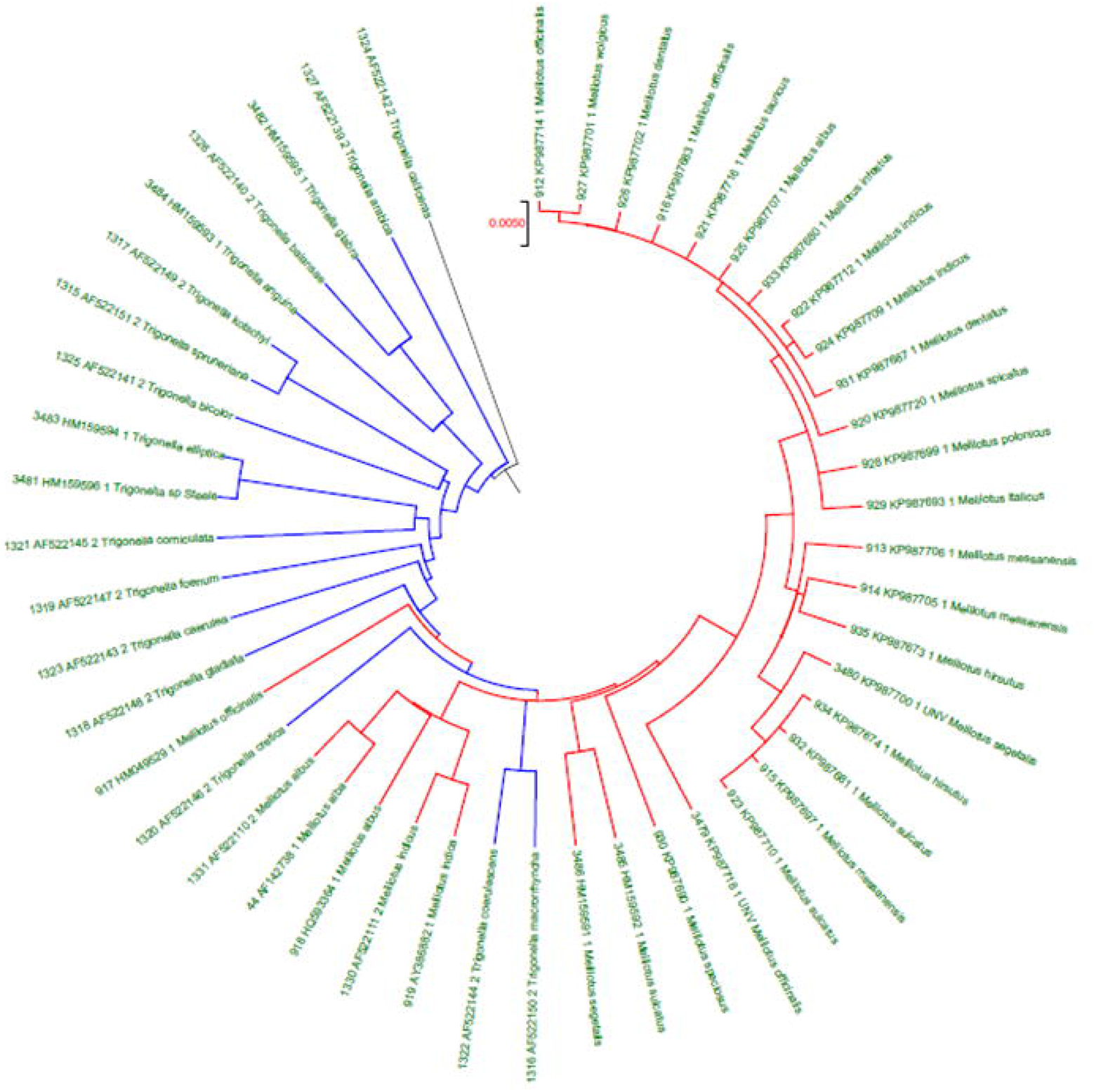
Dalbergia.

**Figure 17.**
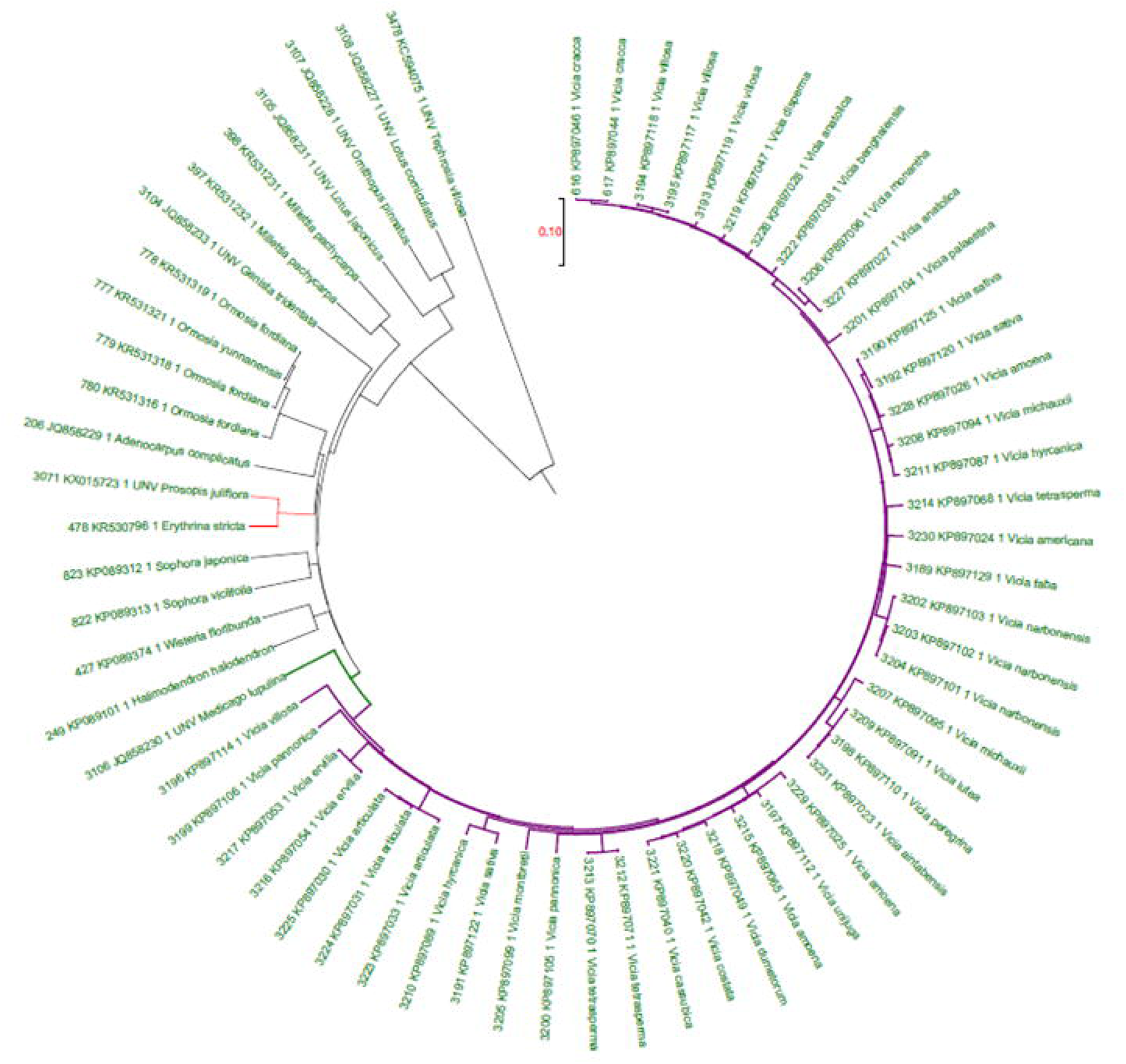
Medicago.

**Figure 18.**
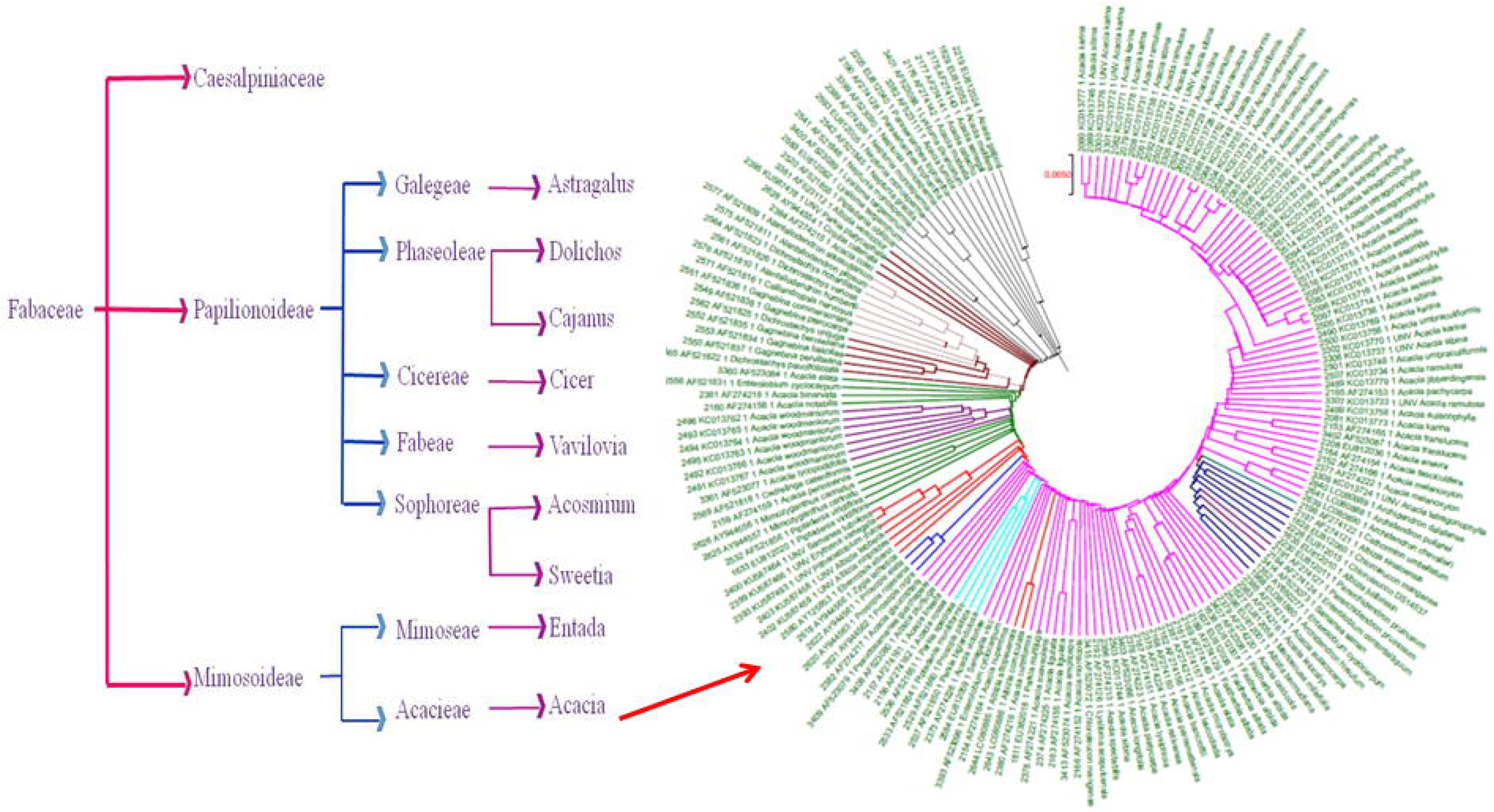
Overview from database.

**Figure 19.**
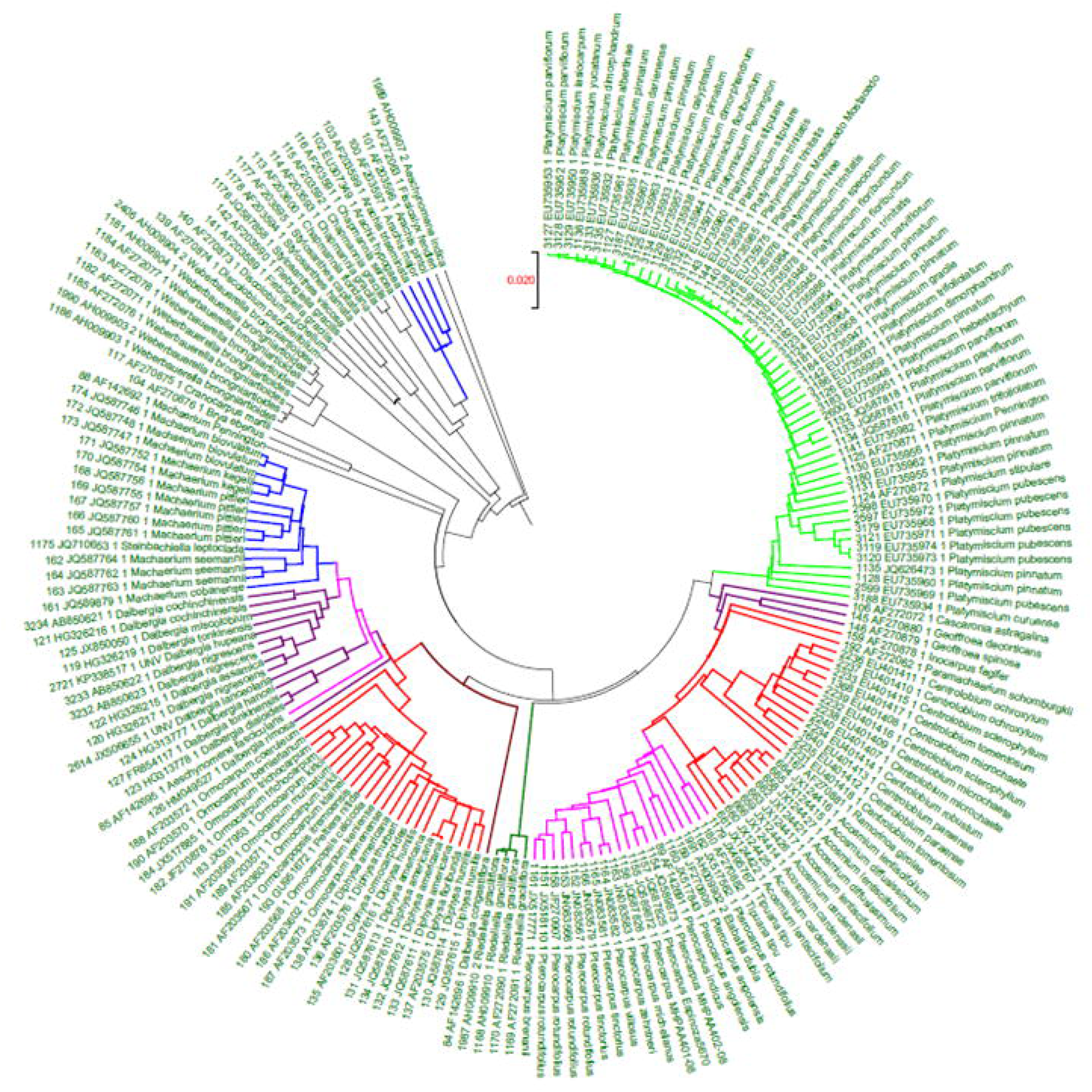
Acosmium group.

**Figure 20.**
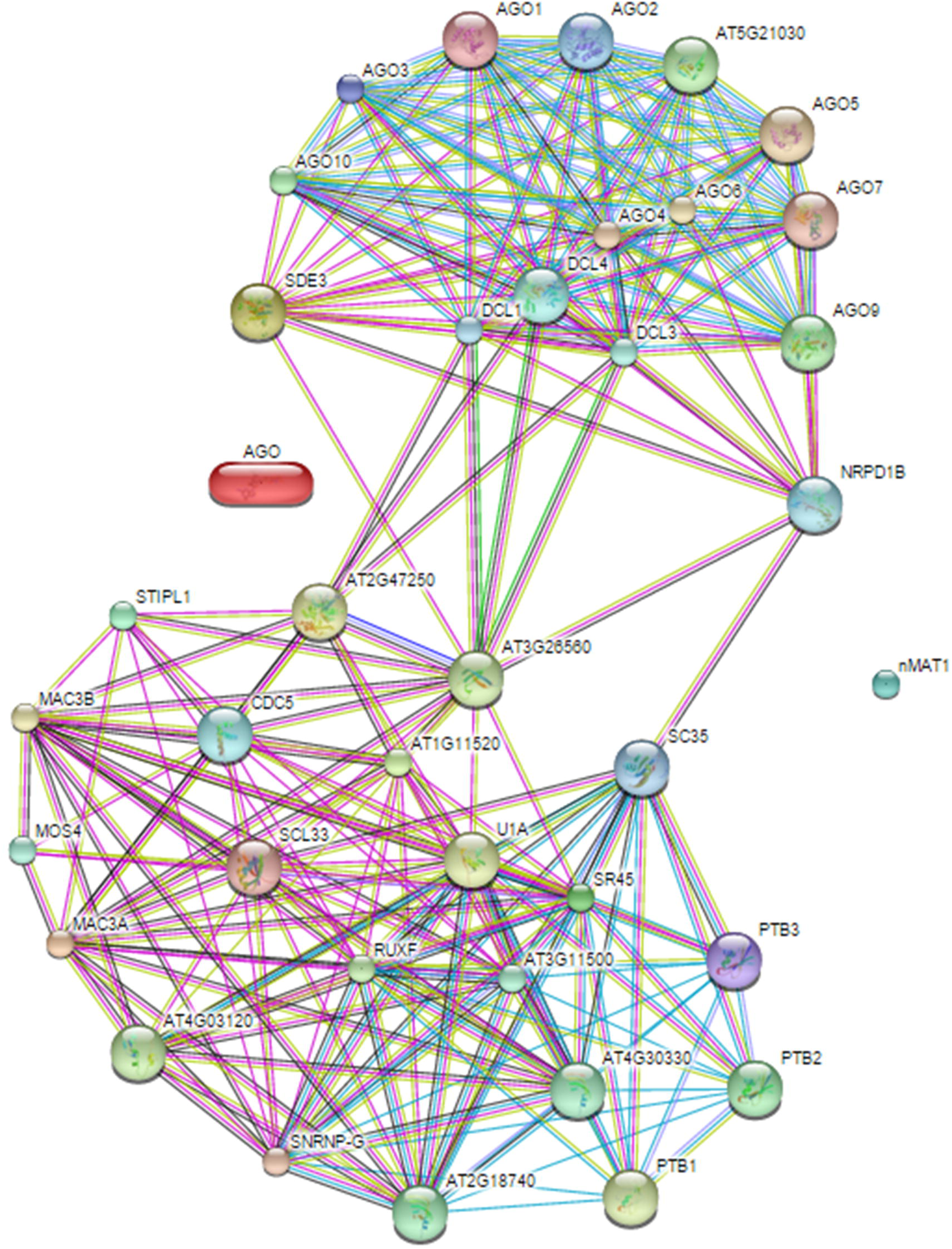
Eriosema group.

Acosmium belongs to sophoreae of papilionoideae. But these sophoreae members were placed along with Dalbergieae members in *matK* based cluster viz., Inocarpus, Centrolobium, Aeschynomene, Dalbergia etc (Fig. 19). Likewise Astragalus is a member of Galegeae. This genus is clustered with members of Erophaca, Swainsona and Clianthus etc.

Cajanus has been placed in a tribe Phaseoleae within a cluster with Eriosema and a member of Rhynchosia but couldn’t find Atylosia here (Fig. 20). It could be possible that phaseoleae is utterly unresolved by *matK*, whereas, Cicer is evidently listed and grouped (Fig. 21).

*Vigna membranacea* could be a close relative of Macrotyloma than other vigna species. Hence, both the genus may be revisited, reconsidered and revised. Vigna forms a different subgroup, whereas Macroptilium is admixed with phaseolus (Fig. 22). Prosopis and Erythrina (Fig. 23); Trifolieae members Trigonella with Melilotus (Fig. 24) are not completely resolved using *matK* sequences.

### In-silico miRNAs targeting *matK*

Of the total reported 8496 miRNAs in viridiplantae only three were (mtr-miR5222, gma-miR1536 and aly-miR3434–5p) found to target maturase sequences. It could be an insight to investigate further detecting the possibilities of AGO complex to migrate inside organelles from cytoplasm effecting gene regulation.

### Combined analysis

Multiple sequence alignment epitomise the inconsistent number of variations in evolutionary stretch in *matK* sequences. Interestingly, the combined *matK* sequences alignment confirms a total of 497 and 251 sites corresponds to variable and parsimony sites respectively with a significant overall mean distance of 0.027 representing 0.38% to 8.85% variants. Our analyses corroborate with previous studies confirming papilionoids and mimosoids as monophyletic groups nested within a paraphyletic Caesalpinioideae excluding Dinizia of tribe Mimoseae.

### Utility and Discussion

We represent FabElm_BarcodeDb, a matK barcode database for scrupulous, brisk and stereotype method for legumes maturaseK identification using short DNA sequence. These barcodes will help in identifying the unknown plant samples, phylogenetic analyses of selected sequences and retrieval of barcode of expected plant species. FabElm_BarcodeDb can be accessed through user friendly web interface (will be made available at http://app.bioelm.com) that provide search options like genus, species, sequence id etc. Such a standardised identification method will be useful for mapping, species identification and sequence analysis. Plant barcodes with additional information remain as a classical and integrative approach for identification of novel species or unknown samples. In earlier reports, surprising sequence discrepancy has directed to re-investigate such morphological /environmental variation, finally results in initial novel taxa detection. As such there is lack of specific database available for legume barcodes. Our exhaustive methodology will be exploited and utilized as a significant database by the scientific community involved in various characteristics of legume barcode with guarantee free access to significant superior datasets. The user friendly database will be having options to add new sequences uploaded in major sequence submission portals.

Plant *matK* is homologous to bacterial maturase (Neuhaus and Link, 1987). Due to high substitution rate at nucleotide as well as amino acids levels, matK shows uncommon evolutionary attributes which indicates its significance. The base substitution rate of matK is three times higher than rbcL (Johnson et al., 1994; Olmstead et al., 1994), denoting matK as a rapidly evolving gene (Soltis and Soltis, 2004). MatK comprises high phylogenetic signal with more parsimony infromation than rbcL and trnT-F (Muller et al., 2006). Furthermore, the molecular information generated from matK has been utilised to decide phylogenetic relationship ranging from trivial to profound taxonomic levels (Johnson and Soltis, 1994; Hilu et al., 2003). Likewise, rearrangement in Adiantum chloroplast genome leads absence of trnK but maintenance of matK (Wolfe et al., 1992). The keep of the matK gene in both species after loss of numerous other genes proposes that matK is expressed and serves a crucial and unique function in plants (Barthet et al., 2007).

Chloroplast gene splicing proteins were already characterized (Khrouchtchova et al., 2012). Amongst these characterised splicing factors, single loci matK is eminent while, other genic factors are encoded in nucleus. InDels are repetitive in matK, nonetheless these InDels predominantly follow a multiple of three pattern, enduring their reading frames. Additionally, the investigation of transition/ transversion proportion in the gene sequences approach are in accordance for both profound as well as superficial taxonomic levels i.e order and subfamily respectively (Liang and Hilu, 1996; Hilu et al., 2003). Alternatively, a number of investigations advocate that matK might be absent/non-functional in few plants due to their swift substitution rate, occasional existence of frame shift InDels and limited circumstances of early stop codons (Kores et al., 2000; Whitten et al., 2000; Kugita et al., 2003; Hidalgo et al., 2004; Jankowiak et al., 2004). Substitutions in matK are comparable with matR and nMat sequences wehre matR represents clear at species level variations, while considering these facts splicing can be directly correlates to speciation.

Phylogenetic reconstruction of Leguminosae is critical to understand the source and divergence of this organically and frugally significant angiosperm. Widespread phylogeny of Leguminosae started with plastid gene, rbcL (Chase et al., 1993; Doyle, 1987; Doyle, 1997) followed by matK gene (Wojciechowski et al., 2004). Additionally, with higher favourable phylogeny evidence of matK compared to rbcL and trnT-F, has manifested itself as it more contributor for parsimony informative characters and considerably additional phylogenetic organization on a typical mean per parsimony-informative site compared to the extremely preserved rbcL (Muller et al., 2006). Furthermore, the molecular evidence iterated through matK is utilised to predict the phylogenetic association from trivial to profound taxonomic stages in several studies (Johnson and Soltis, 1994; Hilu et al., 2003). Likewise, rearrangement in Adiantum chloroplast genome leads absence of trnK nonetheless maintenance of matK (Wolfe et al., 1992). The keep of the matK in species subsequently deficit of a number of genes during evolution recommends matK expression crucial with unique purposes in plants (Barthet et al., 2007).

The phylogenetic exploration of the leguminous plant matK sequences represented in this study achieve our central objective of restructuring a strong molecular phylogeny, along with the fastidious and high resolute resulted the individuality and inter-relationship among numerous clades. Interestingly, the monophyly family, features of associations between outgroups, characteristics and relationships amongst several subgroups are momentarily addressed previously (Luckow et al., 2000, 2003). While, our intensive and combinatorial approach discloses considerable potential of matK sequences in determining key clades representing Fabales with little misplacement may either be due to wrong sampling or sequencing error. This could possibly by lesser efficiency of matK alone to resolve these genus clearly. In figure 6, acacia group the genus Archidendron (sequence id. 2640 and 2641) were grouped within Acacia and scattered. This targeted study evaluating matK gene sequences of numerous legumes indicated inability of maturase to be a solitary barcode biomarker due to lack of enough resources, though it contributes to speciation.

### Conclusion

A detailed investigation of *matK’s* role has direct and indirect implications for plant biology. A study of prospective substrates for matK activity underpins the regulatory role at post-transcriptional photosythetic activities in the presence of light. Only Sauret-Gueto et al. (2006) studied a direct link amongst regulation of plant developmental stage and chloroplast post-transcription processing factors. As such splice variants can lead to set of new proteins altering the phenotype leading to different identity and hence a divergent species. In conclusion, the matK gene with its unique characteristics is an important resource for systematic and evolutionary questions because of its size and high rate of substitutions. But, this single loci may not be sufficient to conclude the genus and species of any legume in few cases. It should be combined with any other barcode loci for species identification. A relatively conserved 39 region and more conserved 59 region gives us two different set of features from matK which can be utilized at a diverse taxonomic hierarchal studies.

The present investigation revealed the need for intense and through study of legume barcodes. Though the role of *matK* is proven to be speciation by splicing, it cannot be considered alone as a barcode loci of legumes.

## Declarations

## Acknowledgements

We are grateful to Dr. Kuldeep Singh, Director, ICAR-NBPGR for providing facilities to conduct this research.

## Availability of data

The datasets processed in this study are bundled and made available with the manuscript as supplementary information and will be available from app.bioelm.com for non-commercial research and education purpose.

## Author contributions

YJK designed and planned the research, BKM, YJK and SC carried out *in-silico* analyses, BKM and YJK developed graphical representations, BKM wrote the initial draft, YJK compiled the results, revised, improved and communicated the manuscript.

## Additional Information

Supplementary information accompanies this paper will be available from online version.

## Consent for publication

Not applicable.

## Ethics approval and consent to participate

Not applicable.

## Figure legends

Figure 21 Cicer group

Figure 22 Macroptilium with phaseolus

Figure 23 Prosopis and Erythrina group

Figure 24 Trigonella with melilotus

Figure 25 Splicing AGO network

Supplementary material 1. Splicing of nMAT medicago

## References

1. Adams KL, Qiu YL, Stoutemyer M, Palmer JD. Punctuated evolution of mitochondrial gene content: high and variable rates of mitochondrial gene loss and transfer to the nucleus during angiosperm evolution. Proceedings of the National Academy of Sciences. 2002;99: 9905–9912.

2. Barthet MM, Hilu KW. Expression of matK: functional and evolutionary implications. American Journal of Botany. 2007;94:1402–1412.

3. Chase MW, Soltis DE, Olmstead RG, Morgan D, Les DH, Mishler BD, Kron KA. Phylogenetics of seed plants: an analysis of nucleotide sequences from the plastid gene rbcL. Annals of the Missouri Botanical Garden. 1993;528–580.

4. Delannoy E, Fujii S, des Francs-Small CC, Brundrett M, Small I. Rampant gene loss in the underground orchid *Rhizanthella gardneri* highlights evolutionary constraints on plastid genomes. Molecular Biology and Evolution. 2011;28:2077–2086.

5. Doyle JJ. Trees within trees: genes and species, molecules and morphology. Systematic Biology. 1997:46:537–553.

6. Doyle JJ. A rapid DNA isolation procedure for small quantities of fresh leaf tissue. Phytochem bull. 1987;19:11–15.

7. Edgar RC. MUSCLE: multiple sequence alignment with high accuracy and high throughput. Nucleic acids research. 2004;32:1792–1797.

8. Funk HT, Berg S, Krupinska K, Maier UG, Krause K. Complete DNA sequences of the plastid genomes of two parasitic flowering plant species, *Cuscuta reflexa* and *Cuscuta gronovii*. BMC Plant Biology. 2007;7:45.

9. Goldman DH, Jansen RK, van den Berg C, Leitch IJ, Fay MF, Chase MW. Molecular and cytological examination of Calopogon (Orchidaceae, Epidendroideae): circumscription, phylogeny, polyploidy, and possible hybrid speciation. American Journal of Botany. 2004;91:707–723.

10. Grewe F, Edger PP, Keren I, Sultan L, Pires JC, Ostersetzer-Biran O, Mower JP. Comparative analysis of 11 Brassicales mitochondrial genomes and the mitochondrial transcriptome of *Brassica oleracea*. Mitochondrion. 2014;19:135–143.

11. Group CPW, Hollingsworth PM, Forrest LL, Spouge JL, Hajibabaei M, Ratnasingham S, Fazekas AJ. A DNA barcode for land plants. Proceedings of the National Academy of Sciences. 2009;106:12794–12797.

12. Hidalgo O, Garnatje T, Susanna A, Mathez J. Phylogeny of Valerianaceae based on matK and ITS markers, with reference to matK individual polymorphism. Annals of Botany. 2004;93:283–293.

13. Hilu KW, Borsch T, Muller K, Soltis DE, Soltis PS, Savolainen V, Chase MW, Powell MP, Alice LA, Evans R, Sauquet H. Angiosperm phylogeny based on matK sequence information. American Journal of Botany. 2003;90:1758–1776.

14. Jankowiak K, Lesicka Joanna, Pacak Andrzej, Rybarczyk A, Szweykowska-Kulinska Z. A comparison of group II introns of plastid tRNALys UUU genes encoding maturase protein. Cell. Mol. Biol. Lett, 2004;9:239–251.

15. Johnson LA, Soltis DE. matK DNA sequences and phylogenetic reconstruction in Saxifragaceae s. str. Systematic Botany. 1994;143–156.

16. Kajita T, Ohashi H, Tateishi Y, Bailey CD, & Doyle JJ. rbcL and legume phylogeny, with particular reference to Phaseoleae, Millettieae, and allies. Systematic Botany. 2001;26:515–536.

17. Khrouchtchova A, Monde RA, Barkan A. A short PPR protein required for the splicing of specific group II introns in angiosperm chloroplasts. Rna. 2012;18:1197–1209.

18. Kores PJ, Weston PH, Molvray M, Chase MW. Phylogenetic relationships within the Diurideae (Orchidaceae): inferences from plastid matK DNA sequences. Monocots: systematics and evolution. 2000;449–456.

19. Kugita M, Kaneko A, Yamamoto Y, Takeya Y, Matsumoto T, Yoshinaga K. The complete nucleotide sequence of the hornwort (Anthocerosformosae) chloroplast genome: insight into the land plants. Nucleic Acids Research. 2003;31:716–721.

20. Sultan LD, Mileshina D, Grewe F, Rolle K, Abudraham S, Głodowicz P, Barciszewski J. The Reverse Transcriptase/RNA Maturase Protein MatR Is Required for the Splicing of Various Group II Introns in Brassicaceae Mitochondria. The Plant Cell. 2016;28:2805–29.

21. Liang H, Hilu KW. Application of the mat K gene sequences to grass systematics. Canadian Journal of Botany. 1996;74:125–134.

22. Mabberley DJ. The plant-book: a portable dictionary of the vascular plants. Cambridge university press.

23. McNeal JR, Kuehl JV, Boore JL, Leebens-Mack J. Parallel loss of plastid introns and their maturase in the genus Cuscuta. PLoS One. 2009;4:e5982.

24. Muller KF, Borsch T, Hilu KW. Phylogenetic utility of rapidly evolving DNA at high taxonomical levels: contrasting matK, trnT-F, and rbcL in basal angiosperms. Molecular phylogenetics and evolution. 2006;41:99–117.

25. Neuhaus H, Link G. The chloroplast tRNA Lys (UUU) gene from mustard *(Sinapis alba)* contains a class II intron potentially coding for a maturase-related polypeptide. Current Genetics,1987;11:251–257.

26. Olmstead RG, Palmer JD. Chloroplast DNA systematics: a review of methods and data analysis. American journal of botany. 1994;205–24.

27. Polhill RM. Classification of the Leguminosae. Phytochemical dictionary of the Leguminosae. 1994;1:35–48.

28. Rundel PW. Ecological success in relation to plant form and function in the woody legumes. Advances in Legume Biology Monogr. Syst. Bot. Missouri Bot. Gard. 1989;29:377–98.

29. Sauret-Gueto S, Botella-Pavia P, Flores-Perez U, Martinez-Garcia JF, San Roman C, Leon P, Rodriguez-Concepcion M. Plastid cues posttranscriptionally regulate the accumulation of key enzymes of the methylerythritol phosphate pathway in Arabidopsis. Plant Physiology. 2006; 141: 75–84.

30. Schmitz-Linneweber C, Williams-Carrier R, Barkan A. RNA immune-precipitation and microarray analysis show a chloroplast pentatricopeptide repeat protein to be associated with the 5′ region of mRNAs whose translation it activates. The Plant Cell. 2005;17:2791–2804.

31. Soltis DE, Soltis PS, Chase MW, Mort ME, Albach DC, Zanis M, & Axtell M. Angiosperm phylogeny inferred from 18S rDNA, rbcL, and atpB sequences. Botanical Journal of the Linnean Society. 2000;133:381–461.

32. Soltis DE, Soltis PS, Morgan DR, Swensen SM, Mullin BC, Dowd JM, Martin PG. Chloroplast gene sequence data suggest a single origin of the predisposition for symbiotic nitrogen fixation in angiosperms. Proceedings of the National Academy of Sciences. 1995;92:2647–51.

33. Soltis DE, Soltis PS. Amborella not a “basal angiosperm”? Not so fast. American journal of botany. 2004;91:997–1001.

34. Sprent JI. Nitrogen acquisition systems in the Leguminosae. In: Sprent JI, McKey D, editors. Advances in Legume Systematics 5. The Nitrogen Factor: Royal Botanic Gardens, Kew; 1994;1–16.

35. Sprent JI. Nodulation in legumes. Royal Botanic Gardens, Kew. 2001.

36. Stern DB, Goldschmidt-Clermont M, Hanson MR. Chloroplast RNA metabolism. Annual review of plant biology. 2010;61:125–55.

37. Turmel M, Otis C, Lemieux C. The chloroplast genome sequence of Chara vulgaris sheds new light into the closest green algal relatives of land plants. Molecular Biology and Evolution. 2006;23:1324–38.

38. Whitten WM, Williams NH, Chase MW. Subtribal and generic relationships of Maxillarieae (Orchidaceae) with emphasis on Stanhopeinae: combined molecular evidence. American Journal of Botany. 2000;87:1842–56.

39. Wolfe KH, Morden CW, Palmer JD. Function and evolution of a minimal plastid genome from a nonphotosynthetic parasitic plant. Proceedings of the National Academy of Sciences. 1992;89:10648–52.

40. Wojciechowski MF, Lavin M, Sanderson MJ. A phylogeny of legumes (Leguminosae) based on analysis of the plastid matK gene resolves many well-supported subclades within the family. American Journal of Botany. 2004;91:1846–62.

41. Steele KP, Vilgalys R. Phylogenetic analyses of Polemoniaceae using nucleotide sequences of the plastid gene matK. Systematic Botany. 1994;126–42.

42. Yasin JK. Smart Way to Nutritional Security in India. Journal of AgriSearch. 2016;3:135.

43. Ramya KT, Fiyaz RA, Yasin JK. SMART agriculture for nutritional security. Current science. 2003;105:1458.

44. Yasin JK. “SMART” to success: not just combinations and permutations. 2015. http://comments.sciencemag.org/content/10.1126/science.1254135.

45. Persson C. Phylogenetic relationships in Polygalaceae based on plastid DNA sequences from the trnL-F region. Taxon. 2001;763–779.

46. Dai X, Zhao PX. psRNATarget: a plant small RNA target analysis server. Nucleic acids research. 2011;39:W155–W159.

47. Kumar S, Stecher G, Tamura K. MEGA7: Molecular Evolutionary Genetics Analysis version 7.0 for bigger datasets. Molecular biology and evolution. 2016;msw054.

48. Luckow Melissa, Miller JT, Murphy DJ, Livshultz Tatyana, A phylogenetic analysis of the Mimosoideae (Leguminosae) based on chloroplast DNA sequence data. Advances in legume systematics. 2003;10,197–220.

49. Luckow M, White PJ, Bruneau A. Relationships among the basal genera of mimosoid legumes. Advances in legume systematics. 2001;9:165–180.

